# Guanine quadruplexes mediate mitochondrial RNA polymerase pausing

**DOI:** 10.1101/2023.10.17.562821

**Authors:** Ryan Snyder, Don Delker, Joshua T Burdick, Vivian G. Cheung, Jason A. Watts

**Affiliations:** Epigenetics and Stem Cell Biology Laboratory, National Institute of Environmental Health Sciences, National Institutes of Health, Research Triangle Park, NC 27709, USA; Integrative Bioinformatics, National Institute of Environmental Health Sciences, National Institutes of Health, Research Triangle Park, NC 27709, USA; Life Sciences Institute, University of Michigan, Ann Arbor, MI 48109, USA; Department of Pediatrics, Division of Neurology, University of Michigan, Ann Arbor, MI,USA; Department of Medicine, University of Michigan, Ann Arbor, MI 48109, USA

**Author notes:** Please send correspondence to JAW.

## Abstract

The information content within nucleic acids extends beyond the primary sequence to include secondary structures with functional roles in cells. Guanine-rich sequences form structures called guanine quadruplexes (G4) that result from non-canonical base pairing between guanine residues. These stable structures are enriched in gene promoters and have been correlated with the locations of RNA polymerase II pausing (Pol II). While promoter-proximal RNA polymerase pausing regulates gene expression, the effects of guanine quadruplexes on gene transcription have been less clear. We determined the pattern of mitochondrial RNA polymerase (mtRNAP) pausing in human fibroblasts and found that it pauses over 400 times on the mitochondrial genome. We identified quadruplexes as a mediator of mtRNAP pausing and show that stabilization of quadruplexes impeded transcription by mtRNAP. Gene products encoded by the mitochondrial genome are required for oxidative phosphorylation and the decreased transcription by mtRNAP resulted in lower expression of mitochondrial genes and significantly reduced ATP generation. Energy from mitochondria is essential for transport function in renal epithelia, and impeded mitochondrial transcription inhibits transport function in renal proximal tubule cells. These results link formation of guanine quadruplex structures to regulation of mtRNAP elongation and mitochondrial function.

## Introduction

Mitochondria are organelles with diverse functions in cellular homeostasis, including energy production in the form of ATP. They are unique among organelles in human cells by having their own genome which is transcribed by the mitochondrial RNA polymerase. Dysregulation of mitochondrial gene expression contributes to several health conditions including cancer, cardiovascular and kidney diseases. Thus, an understanding of mitochondrial transcription is vital to understanding mitochondrial dysfunction in disease.

The mitochondrial genome is a double-stranded circular DNA of 16.5 kb that is not associated with histones. It encodes 13 proteins which are essential components of the electron transport chain, as well as 2 rRNA and 22 tRNAs which are necessary for translation of the mitochondrial mRNA(1).

Transcription by the mitochondrial RNA polymerase (mtRNAP) initiates from two major promoters, the light strand promoter (LSP) and the heavy strand promoter (HSP_1_), and results in the synthesis of two nearly genome-length polycistronic transcripts(2). These transcripts, which do not contain introns, are then processed to release the mature mRNAs(3). The proteins needed to initiate transcription have been defined and reconstituted in vitro(2,4), yet the pattern by which the mtRNAP synthesizes the polycistronic RNA has been less well characterized.

In the nucleus, transcription by RNA polymerase II (Pol II) is discontinuous. It transcribes the DNA in a series of pause and elongation steps(5). These pauses are determinants of transcript abundance and therefore gene expression. Pol II pausing in gene promoters has been correlated with guanine-rich sequences where G-quadruplexes (G4s) form(6). G4s are three-dimensional structures that arise from non-canonical Hoogsteen base pairing between runs of guanines, leading to the formation of G-quartets which stack onto one another to form a guanine quadruplex(7). In vitro, guanine quadruplexes are sufficient to pause Pol II(8). While quadruplexes are associated with paused Pol II, there is conflicting evidence regarding the effect of quadruplexes on transcription. Stabilization of guanine quadruplexes can repress gene expression(9,10), yet genes with guanine quadruplexes in the promoter-proximal-region have higher average expression(11,12).

mtDNA is GC rich and has among the highest GC content when compared to chromosomes in the nucleus(13). The two strands have a different number of guanines, giving rise to a guanine-rich RNA transcript and a guanine-poor RNA transcript. Guanine quadruplex formation downstream of the light strand promoter has a role in the switch between mitochondrial transcription or replication, demonstrating that these structures function in transcription regulation(14,15). However, a regulatory function for quadruplex formation in the transcription of the body of the polycistronic genes has not been demonstrated.

We have been interested in the role of nucleic acid structure in the regulation of transcription. Recently, mtRNAP was shown to pause near the light strand promoter(16). In this study, we determined the pattern of pausing of the mitochondrial RNA polymerase as it synthesizes RNA in human fibroblasts. We found over 400 sites where it pauses and identified guanine quadruplexes as a key determinant of mtRNAP pausing. Here, we describe a role for guanine quadruplexes in the regulation of mitochondrial gene expression and mitochondrial function.

## Methods

### Circular dichroism

DNA and RNA oligonucleotides (Supplemental Table 1) were diluted to 10 μM in TE buffer supplemented with 100 mM NaCl, then heated to 85°C for 2 minutes and allowed to cool slowly to room temperature. The samples were then loaded into a 1-mm path quartz cuvette and analyzed under N_2_ in a J-1500 spectrophotometer (Jasco). The CD spectra were collected at 25° C from 220-320 nm with 1-nm bandwidth and 0.1-nm pitch.

### DNA-IP

Fibroblasts cells cultured in 1 μM RHPS4 or DMSO were cross-linked for 10 minutes in 1% formaldehyde and processed as for chromatin IP(17). G-quadruplexes were immunoprecipitated with 5 μg 1H6 antibody (Millipore) or 5 μg rabbit IgG (Sigma) and fold enrichment determined by qPCR. Primers are listed in Supplemental Table 1.

### Extracellular flux analysis

For Seahorse analysis (XFe24, Agilent Technologies), fibroblasts were seeded at a density of 20,000 cells/well in XFe24 cell culture microplates. Fibroblasts were treated with 1 μM RHPS4 or DMSO for 2 and 22 hours, then serum-starved (in medium containing treatment but no serum), for an additional 2 hours. Following treatment, cells were washed twice with serum-free XF DMEM medium (Agilent Technologies, supplemented with 100 μM pyruvate, 200 μM L-glutamine, and 1.1 μM glucose), then allowed to incubate in XF DMEM for 1 hour at 37ºC and 0% CO_2_. Cells were tested for oxygen consumption rate and ATP production following the manufacturer’s instructions for the Seahorse XF Cell Mito Stress Test Kit (Agilent Technologies). Measurements were taken both at baseline prior to mitochondrial stress and following the addition of 2 μM FCCP uncoupling agent.

### Fibroblast Cell culture

Foreskin fibroblasts from healthy newborns (Penn SBDRC) were cultured at 37°C with 5% CO2 in MEM medium (Thermo-Fisher) supplemented with 10% fetal bovine serum, 1% L-glutamine, and 1% penicillin-streptomycin. Cells were passaged every 72 hours using Trypsin-EDTA (0.05%). Culture media was supplemented with 1 μM RHPS4 (Fisher) or equal volume DMSO for 24 hours. Cells were then harvested for PRO-seq, DNA-IP, and gene expression analysis.

### Guanine quadruplex forming sequence

Guadruplex(G4)-forming sequences were identified using QGRS (Kikin et al., 2006) with a spacer of 1-7 nucleotides between each run of consecutive guanines, or using G4Hunter (Brázda et al., 2019) with default settings. The G4-forming sequence scores at sites with or without paused mtRNAP were plotted. We downloaded G4 ChIP-seq (GEO accession GSE107690) (Mao et al., 2018; Zhang et al., 2021) and we extracted information on the G-quadruplexes at the 465 pause sites and random set of sites without mtRNAP pausing, those data are shown in Figures 2C.

**Figure 1.**
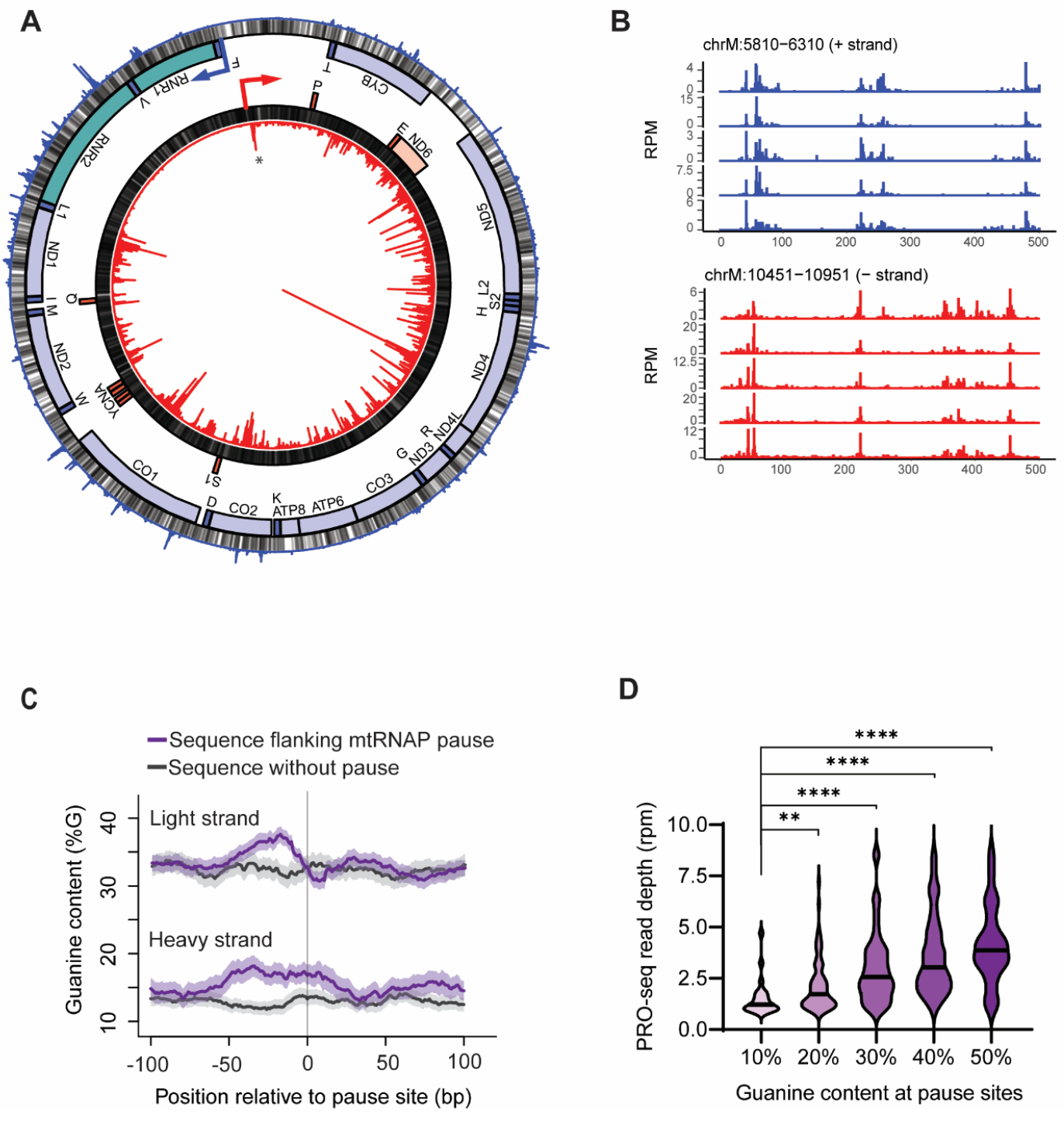
mtRNAP pause throughout the mitochondrial genome. (A) Locations where mtRNAP pauses along the mitochondrial genome. From the center: PRO-seq signal from the light strand shown as a circularized bar graph (red), guanines (grey), genes on the light strand (mRNA red, tRNA dark red), genes on the heavy strand (rRNA green, mRNA blue, tRNA dark blue), guanines (grey), PRO-seq signal from the heavy strand shown in the outer circularized bar graph in blue. The location of a previously identified promoter-proximal pause is indicated (*). Y-axis is 0 to 50 RPM for both heavy and light strands. Data are average PRO-seq signal from primary fibroblasts (N=5). (B) Guanine content is significantly higher at mtRNAP pause sites (G-rich stand P<10^−13^, G-poor strand P<3×10^7^, Wilcoxon test). (C) Violin plots of mtRNAP abundance by guanine content upstream of the pause (*P<0.05, ****P<0.0001; Mann-Whitney U-test).

**Figure 2.**
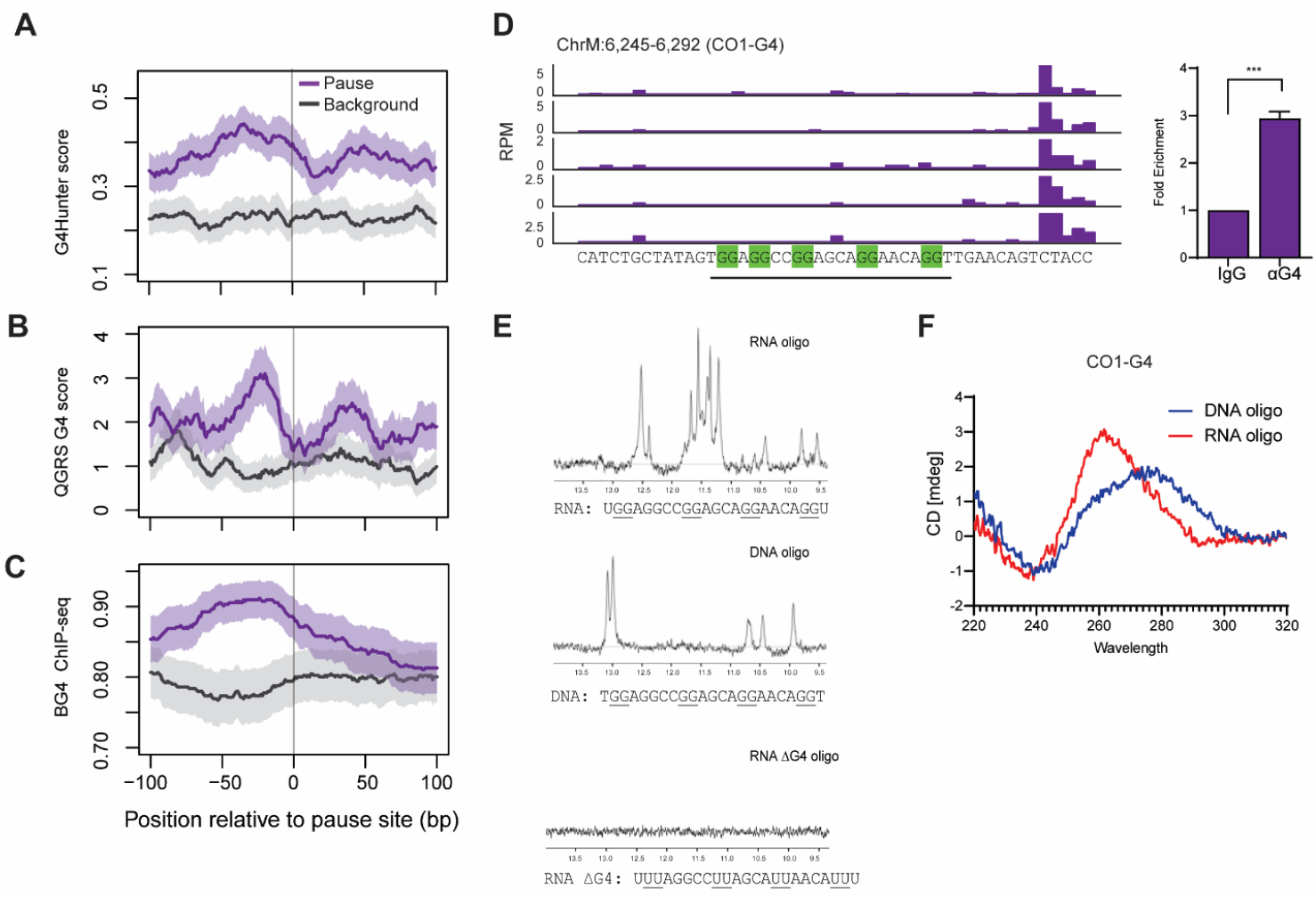
G4 are enriched upstream of mtRNAP pause sites. (A,B) G-quadruplexes were quantified by two algorithms, (A) G4Hunter (B) QGRS, and the abundance of G-quadruplexes for the 100 nucleotides up- and downstream of the 465 mtRNAP pause sites are shown in the Y-axis as the scores (error bands represent S.E.M.). (C) The abundance of G-quadruplexes determined experimentally by BG4 ChIP-seq are also shown for 100 nucleotides up and downstream of the 465 mtRNAP pause sites (error bands represent S.E.M.), (D) G4 forming sequence (underlined) present upstream of pause site in CO1 gene. Corresponding DNA-IP shows enrichment of G4 where mtRNAP pauses (N=4; ***P<0.01; t-test). (E) NMR spectra for CO1-G4 sequences in RNA, or DNA or where the guanines are converted to uracil to prevent G4 formation. NMR chemical shifts in the range of 10-12 ppm indicate Hoogsteen base-pairing indicative of quadruplex formation. (F) Circular dichroism spectra of RNA (red) or DNA (blue) oligos corresponding to sequences in panel D. The pattern with minima at 240 nm and maxima at 260 nm is consistent with an RNA G-quadruplex.

### Gene expression analysis

mRNA expression was examined with the NanoString platform (www.nanostring.com) utilizing a custom mitochondrial codeset (MG_NIH_C10576) that measures 174 endogenous and 8 housekeeping (HK) RNAs(18). 25 ng of each total RNA sample was prepared as per the manufacturer’s instructions. RNA expression was quantified on the nCounter Digital Analyzer™, and data adjusted with positive and negative assay controls as well as highly correlated housekeeping genes were generated with nSolver (v4.0)™ software. Digital gene expression values from cells treated with RHPS4 were normalized to expression in vehicle treated cells and plotted in PRISM.

### Immunoblot

Translation of mitochondrially-encoded protein MT-CO1, nuclear-encoded TOMM20, and phosphorylation of AMPK were assessed by western blotting. Total proteins were extracted from RPTEC transwell cultures using RIPA buffer supplemented with protease inhibitor. Protein yields were verified by BCA assay (Pierce). Samples were denatured in LDS sample buffer and Reducing Agent (Invitrogen/Thermo-Fisher) at 70°C for 10 min and then electrophoretically separated by SDS-PAGE in 4-12% Bis-Tris precast gels in MOPS buffer (Invitrogen/Thermo-Fisher). Gels were transferred to PVDF membranes using the iBlot2 semi-dry transfer device (Invitrogen/Thermo-Fisher) and washed with PBS plus 0.1% Tween 20 (PBST). Blots were then blocked at room temperature for 1 hour with 5% bovine serum albumin (BSA) in PBST, then primary antibodies were applied at 4°C on a shaker overnight.

Primary antibodies were diluted 1:1000 in blocking buffer and included rabbit α-TOMM20 (clone D8T4N, Cell Signaling Technologies #4240), mouse α-MT-CO1 (clone 1D6E1A8, AbCam #ab14707), rabbit α-phospho-AMPK (Thr172)(clone 40H9, Cell Signaling Technologies #2535), and rabbit α-GAPDH (pAb, AbCam #ab9485) as a loading control. Blots were washed with PBST and then incubated with HRP-conjugated secondary antibodies at 1:5000 dilution in wash buffer for 1 hour at room temperature. Secondary antibodies were either Goat α-Rabbit IgG H&L HRP (AbCam, #ab205718) or Goat α-Mouse IgG H&L HRP (AbCam, #ab205719). Blots were washed again and imaged using enhanced chemiluminescent reagent (ECL, Cytiva) on an Azure Sapphire Biomolecular Imager. Densitometry was calculated using AzureSpot software following “rolling ball” background subtraction and results were normalized to GAPDH expression for each target.

### Microscopy

Human primary skin fibroblasts were cultured onto 35mm collagen-coated petri dishes with a No. 1.5 glass coverslip bottom (Mat Tek) at 10,000 cells/cm^2^. Cultures were allowed to reach 80% confluence and treated with either 1 μM RHPS4 or DMSO in complete medium for 2 hours or 22 hours, then treated in serum-free MEM for an additional 2 hours (for the 4-hour and 24-hour treatments, respectively). Mitochondrial staining was accomplished using the live-cell stain MitoTracker Red CMXRos dye (Invitrogen). Cells were stained with 500 nM MitoTracker during their 2-hour serum-starvation, as well as 1 μg/mL DAPI counterstain. Just prior to imaging, media and dyes were aspirated and cells were washed twice with sterile PBS. A thin layer of 100 μL phenol red-free basal medium was used to keep cells hydrated and cultures were enclosed in a 5% CO_2_ chamber during imaging. Imaging of live stained fibroblasts was performed using the Dragonfly spinning-disk confocal microscope (Andor), which was necessary to quickly capture RHPS4 fluorescence without photobleaching. Images were captured at 40x magnification.

To image guanine quadruplex abundance in fibroblasts, cells were grown and treated as described above, then washed and fixed for 20 minutes in 4% paraformaldehyde in PBS. Fixed cells were then permeabilized using 0.5% Triton-X 100 in wash buffer (0.1% Tween 20 in PBS) for 20 minutes. Cells were blocked for 1 hour at room temperature with 1x Fish Gelatin Blocking Agent (Biotium, #22010) in wash buffer. A mouse-derived monoclonal antibody targeting DNA G-Quadruplexes (clone 1H6, Millipore #MAE1126) was diluted 1:200 in blocking buffer and incubated with the cells overnight at 4°C on an orbital shaker. Cultures were then washed 5x 5 minutes each and incubated with Alexa594-conjugated Goat α-Mouse IgG H&L (Abcam, #ab150116) diluted 1:500 in wash buffer for 1 hour at room temperature, then washed and counterstained with DAPI. Imaging of fixed cells was performed using the Cytation 5 Multi-Mode Reader (BioTek). Total G4 staining was quantified by applying a digital threshold of intensity for each image using ImageJ software, generating binary images which were then pixel-counted for percent of area coverage.

### Nuclear Magnetic Resonance Spectroscopy

Oligos were dissolved in PBS buffer including 10% ^2^H_2_O and 20 μM trimethylsilylpropanesulfonic acid (DSS). Oligos were further annealed by incubating at 95 C and allowed to slow cool to room temperature in a heat block. 1D-NMR spectra were acquired at 5C on an 800 MHz Agilent DD2 NMR spectrometer with a cryogenically cooled probe. The oligos used for CD and NMR spectroscopy were listed in Supplemental table 1.

### PRO-seq

PRO-seq data from fibroblasts from five healthy individuals were downloaded from(19) and reads were aligned to the GRCh37 (hg19) build of the human genome using GSNAP(20) (version 2013-10-28) with the mitochondrial genome treated as a circle. BAM files were generated, and data normalized to reads per million mapped reads (RPM). To define pauses, we transformed the depth of PRO-seq read-ends at each base to a Z-score relative to the mean and standard deviation of read-end depth centered in a 201 base window using the rtracklayer R package(21) and custom R scripts. Base positions that had a Z-score ≥3 and at least 1 RPM coverage were called peaks. Locations with a peak of mtRNAP occupancy in two or more samples were considered as pause sites. We tested a range of Z-score thresholds from 3 to 6, and using higher thresholds did not substantively change the conclusions from downstream analysis (see Supplemental Figure 1C).

To test is locations with paused mtRNAP are associated with mitochondrial diseases, a list of mitochondrial SNPs was downloaded from MitoMap https://www.mitomap.org/MITOMAP. Bed files were created from our 465 mitochondrial pause sites and 540 mitochondrial disease associated SNPs. The bedtools intersect tool from the bedtools suite https://bedtools.readthedocs.io/en/latest/index.html was used to determine overlaps between disease SNPs and pause sites. 26 SNPs within mitochondrial pause sites (approx. 6% of sites) were identified.

To measure nascent transcription in cells after stabilization of guanine quadruplexes, human skin fibroblasts were treated with 1 μM RHPS4 (dissolved in DMSO) for 24 hours and PRO-seq from foreskin fibroblasts was performed as described above with the following modifications. The 3’ ligation adapter used oligo 5’p-NNNNNNGAUCGUCGGACUGUAGAACUCUGAAC-/3InvdT/, which contains a unique molecular index (UMI). PRO-seq was sequenced on an Illumina HiSeq 2500 instrument to a depth of >150 million reads per sample. Sequence reads were trimmed and aligned to hg19. Identical reads sharing a UMI were considered duplicates and removed. The 3’ ends of reads mapping to the mitochondrial genome were determined and normalized to the total number of reads mapping to the nuclear genome.

### Transporter Assay

To assess the functional consequences of mitochondrial polymerase pausing, hTERT-immortalized renal proximal tubule epithelial cells (RPTECs) (ATCC, #CRL-4031) maintained in serum free media were seeded on the bottom of collagen-coated permeable transwell membranes (EMD Millipore, #PICM01250) and allowed to form epithelial monolayers. The integrity of the cellular barrier was monitored daily by transepithelial electrical resistance (TEER) across the permeable membrane using a Millicell ESR-2 Voltohmmeter (EMD Millipore) and was confirmed to have increased over 30 days in culture (Supplemental Figure 4A). Formation of intact epithelium was verified by measuring diffusion of Antonia Red Dextran 20 (Sigma, #18650) across the membrane and epithelial cultures which prevented dextran diffusion were used for subsequent analysis. To test the effect of loss and rescue of ATP generation by mitochondria, RTPEC epithelial culture were treated with either 1 μM RHPS4, 1 μM RHPS4 plus 1 mM ATP, or DMSO vehicle for 24 hours, then a fluorescently-labeled glucose analogue (2-NBDG, Sigma #72987) was added to the lower chamber of the transwell insert. Fluorescence in the upper chamber was measured at 499ex/520em after 72 hours using the Cytation 5 Multi-Mode Reader (BioTek).

## Results

### mtRNAP pauses frequently during transcription

In primary human skin fibroblasts from five individuals, we use the precision nuclear run-on assay (PRO-seq) to isolate nascent RNA with actively transcribing RNA polymerase in their 3’ end. We then sequenced the nascent RNA and mapped the locations of the associated mtRNAP on the mitochondria genome to single-nucleotide resolution.(5) (19) The small size of the mitochondrial genome allows us to sequence the nascent RNA deeply, to an average of 250-fold coverage in each individual. As in previous studies, we defined pausing by where the mitochondrial RNA polymerase accumulates (see Methods)(22). Prior to this study there was only one site where the mitochondrial RNA polymerase was known to pause, here we found 465 mtRNAP pause sites (Figure 1A), including the known pause location near the light strand promoter (Supplemental Figure 1A). This large number of mtRNAP pause sites allow us to assess what mediates the polymerase pausing. In contrast to the Pol II that pauses at the gene promoters, mtRNAP does not preferentially pause at the 5’ or 3’ ends of the mRNA nor tRNA genes (Supplemental Figure 1B), instead we found mtRNAP pauses throughout both polycistronic transcripts. Figure 1B shows mtRNAP pausing in all five individuals profiled and the locations are highly correlated between individuals. The mtRNAP pauses in both rRNA genes, in 12 of 13 mRNA genes, and 5 of the 22 tRNA genes (Figure 1A). Comparison of pause site locations with MitoMap showed that 26 (6%) of the pause sites overlap with disease associated sequence polymorphisms. Taken together, we find that mtRNAP pauses frequently in the mitochondrial genome both in the promoter and in the gene body.

### Guanine Quadruplexes are upstream of locations where mtRNAP pauses

A distinctive feature of the mitochondria genome is that one strand is G-rich (31% guanine) and the other is G-poor (13% guanine; see Figure 1A grey bars). Comparison of where the mtRNAP pauses between the two strands shows that the mtRNAP pauses significantly more often (P<0.0001; Chi-square) on the G-rich strand as compared to the G-poor strand (Figure 1A). While there are more pauses on the G-rich strand (N=314 G-rich versus N=151 G-poor), paused mtRNAP is associated with guanine-rich sequences on both strands. Guanines are significantly enriched upstream (5’) of where the mtRNAP pauses (G-rich stand P<10^−13^, G-poor strand P<3×10^7^, Wilcoxon test; Figure 1C and Supplemental 1C). Further, when we grouped sites by the guanine content in the 50 nucleotides upstream of the pause site, we found more guanines were associated with significantly more paused mtRNAP (P<0.0001, ANOVA; Figure 1D).

Given that guanine-rich sequences have a propensity to form secondary structures such as G-quadruplexes (G4 as coined by Sen & Gilbert)(7), we asked if these secondary structures promote mtRNAP pausing. We used G4-hunter and QGRS mapper,(23,24) to identify G4-sequences and determine their locations relative to mtRNAP pause sites. Data from both algorithms showed that G4 sequences with doublet guanine repeats such as G_2_X_1-7_G_2_X_1-7_G_2_ are enriched about 20 to 40 bases upstream of mtRNAP pause sites (Figure 2A, 2B). The BG4 antibody recognizes G4 and has been used for G4-immunoprecipitation followed by sequencing from human cells(12). Analysis of BG4 ChIP-seq further confirms that quadruplexes form in cells and are enriched upstream of mtRNAP pause sites (Figure 2C). Together, the analyses show that the G-rich sequences that can form G-quadruplex structures are found upstream of paused mtRNAP.

### G-quadruplexes form in nascent RNA

Two scenarios can explain G4 formation upstream of RNA polymerase pause sites, the non-template DNA and the newly synthesized RNA can form G-quadruplexes to mediate the pausing. To decipher if one or both of these occur, we directly assessed the propensity of the non-template DNA sequences and their corresponding RNA sequences to form G-quadruplex structures. We selected a pause site in the *MT-CO1* gene with a predicted G4 forming sequence (CO1-G4). Figure 2D shows in all 5 fibroblast samples the mtRNAP pauses downstream of the CO1-G4 sequence which follows the G_2_X_1-7_G_2_X_1-7_G_2_X_1-7_G_2_ pattern. We performed G4-immunoprecipitation in fibroblasts and found that G4s are significantly (P<0.01; t-test) enriched at the CO1-G4 sequence (Figure 2D, right). We synthesized oligonucleotides for the DNA and RNA sequences that correspond to CO1-G4 and then we assessed their secondary structure using nuclear magnetic resonance (NMR) spectroscopy and circular dichroism (CD). In nuclear magnetic resonance spectroscopy, Hoogsteen base-pairing between guanine residues protects the imino hydrogen giving rise to characteristic chemical shifts in the range of 10-12 ppm, whereas Watson-Crick base pairs gives shifts in the range of 13-14 ppm. Figure 2E shows the NMR spectroscopy analysis which indicates that guanine quadruplexes formed in the CO1 RNA but not the DNA. The RNA oligonucleotides gave the characteristic pattern of guanine quadruplexes with chemical shifts in the range of 10-12 ppm, whereas these shifts were absent in both the DNA oligonucleotide as well as an RNA oligonucleotide where the guanines were replaced with uridine. To further characterize the secondary structure of the CO1-G4, we performed circular dichroism spectroscopy. We found the RNA oligo gave the characteristic spectra of parallel G-quadruplexes with minima at 240 nM and maxima near 260 nM; whereas, the CD spectra for the DNA oligo showed an unfolded molecule (Figure 2F), consistent with the results from NMR spectroscopy.

We extended this analysis to the guanine quadruplex forming sequences in the genes including *ND6* and *CYB*. We found the RNA oligos exhibited spectra consistent with parallel quadruplexes, whereas the corresponding DNA oligos showed a mix of unfolded molecules, parallel or antiparallel quadruplexes (Supplemental Figure 2). This shows G4 can form in both DNA and RNA, but sequences with shorter dinucleotide guanine repeats preferentially form quadruplexes in RNA.

### Guanine Quadruplexes pause mtRNAP transcription

Having found that the guanine-rich DNA and nascent RNA sequences upstream of the mtRNAP pause sites form quadruplex structures, we asked if the secondary structure is indeed needed to mediate pausing. We treated skin fibroblasts with RHPS4(25), a small molecule that localizes to the mitochondria and stabilizes guanine quadruplex structures(26). Using fluorescence microscopy, we confirmed that RHPS4 accumulates in the mitochondria by 24 hours (Figure 3A) and leads to greater abundance of guanine quadruplexes (Figure 3B). We performed PRO-seq to assess the pattern of mtRNAP pausing when guanine quadruplexes are stabilized. As the mtRNAP synthesizes nascent RNA, they form quadruplexes and in the presence of RHPS4 the quadruplexes are stabilized and result in mtRNAP pausing at the 5’ end of the polycistronic transcripts. Due to the mRNAP pausing in the proximal region there are fewer mtRNAP at the 3’ end of the polycistronic transcript as shown from replicate PRO-seq samples in Figure 3C.

**Figure 3.**
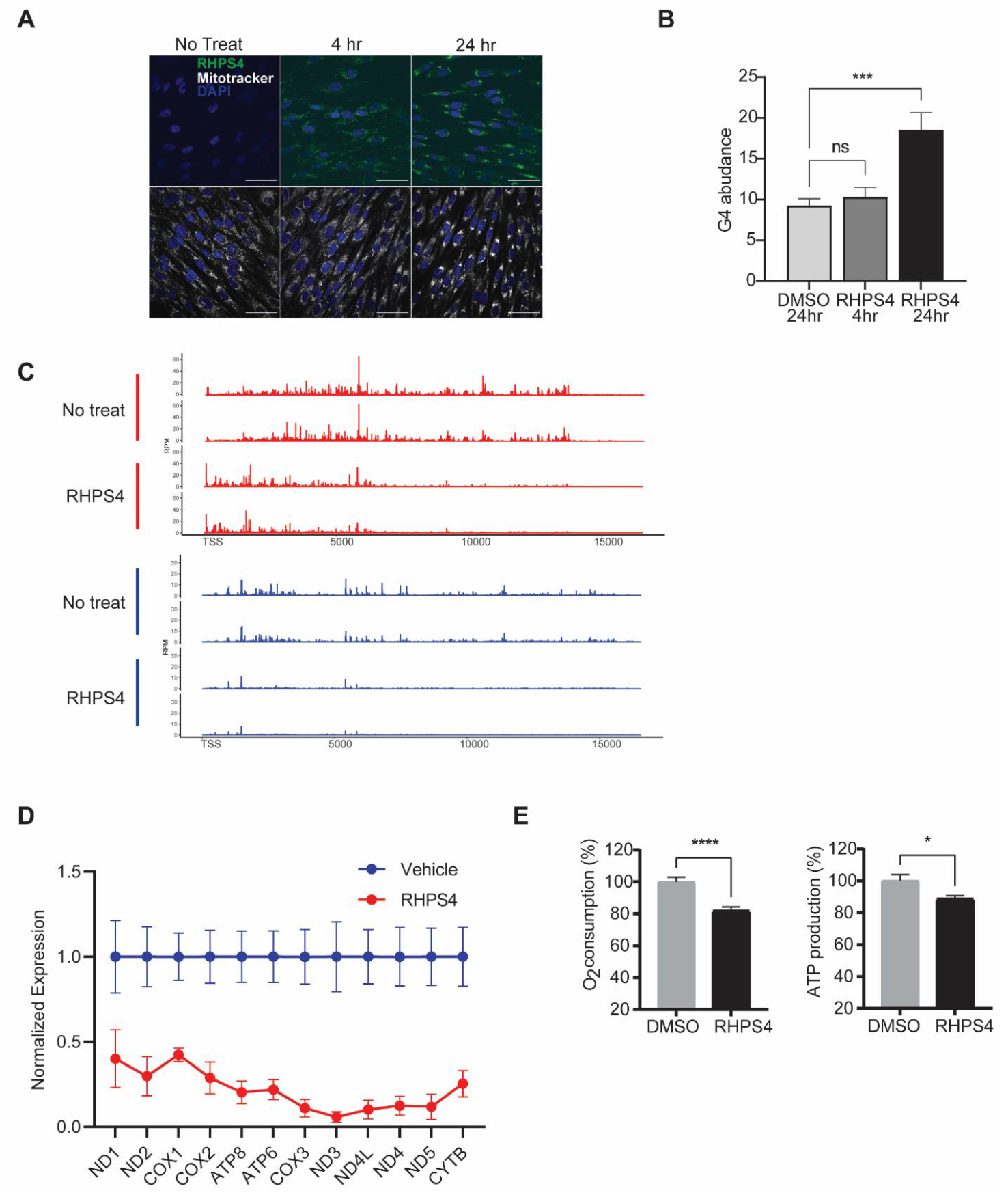
Stable G4 lead to more paused mtRNAP. (A) Confocal image showing RHPS4 (green) localization in the cytoplasm following 24 hrs of drug treatment. Scale bar is 50 μm. (B) Bulk G4 abundance measured by immunofluorescence, expressed as percentage of intracellular area labeled, before and after RHSP4 treatment. (C) PRO-seq data from biologic replicate experiments showing mtRNAP distribution before and after treatment with RHPS4. Y-axis is RPM. RHPS4 results in a shift in mtRNAP localization with more polymerase at the 5’ end of the polycistron and decreased mtRNAP at the 3’ end of polycistron. G-rich light strand transcripts (red). G-poor heavy strand transcripts (blue) (D) Mitochondrial gene expression before and 24 hours after RHPS4 treatment normalized to vehicle expression in arbitrary units (N=3; P<0.0001, one-sided ANOVA, error bars=S.E.M.). (E) Basal oxygen consumption and ATP production before and after 24 hours of RHPS4 treatment are shown. (N=6; *P<0.05, ***P<0.001, t-test, error bars=S.E.M.).

### Guanine quadruplex pausing regulates mitochondrial function

To further examine the effect of the G-rich sequences and G4 structures, we turned to more functional readouts of the effect of RHPS4. We measured the expression of mitochondrially-encoded genes in primary skin fibroblasts after treatment with RHPS4. We found stabilization of guanine quadruplexes lead to a significant decrease in the expression of the 12 polycistronic mRNAs encoded on the heavy strand, including *MT-CO1* (Figure 3D). There is a significant correlation (r=0.53; P<0.05) between gene expression level and the distance of the polycistronic gene from the start of transcription, where genes further from the TSS had lower expression. This is consistent with the pattern of mtRNAP transcription measured by PRO-seq, where there is less polymerase at the distal end of the polycistronic gene.

The proteins encoded in the mtDNA are components of protein complexes in the electron transport chain. For example, *CO1* encodes the cytochrome c oxidase 1 and is a component of complex IV in the electron transport chain. Given the requirement for mitochondrial gene products for cellular respiration, we measured the oxygen consumption rate and ATP production in skin fibroblasts treated with RHPS4. We found that stabilization of G4 leads to significantly lower basal O_2_ consumption and lower ATP generation (Figure 3E). Thus, under conditions that increase mtRNAP pausing, mitochondrial energy production was also impaired.

We repeated the above gene expression analysis in renal proximal tubule cells since mitochondrial function is essential for their solute transport function during urine production. The kidney is a highly metabolically active, with consumption equivalent to that of the heart(27). Like primary fibroblasts, treatment with RHPS4 for 24 hours resulted in accumulation of RHPS4 in the mitochondria and a distance dependent decrease in the expression of mitochondrial transcripts on the heavy strand, including *MT-CO1* (Supplemental Figure 3A,B). To investigate whether altered mtRNAP pausing can alter cellular function, we turned to a transporter assay in renal proximal tubule cells. We seeded cells on a transwell insert and allowed to form an epithelial layer (Figure 4A; Supplemental Figure A-C). We treated cells with RHPS4 and then measured protein expression by immunoblotting. We found no significant differences in the abundance of the nuclear encoded mitochondrial membrane protein TOMM20 but saw decreased expression of the mitochondria encoded CO1 (Figure 4B). This indicates there is no appreciable change in mitochondria abundance in the presence of RHSP4, yet transcription by mtRNAP is impaired. Following treatment with RHPS4 there was a decrease in the energy production resulting in ATP stress, evidenced by increased AMPK phosphorylation (Figure 4B).

**Figure 4.**
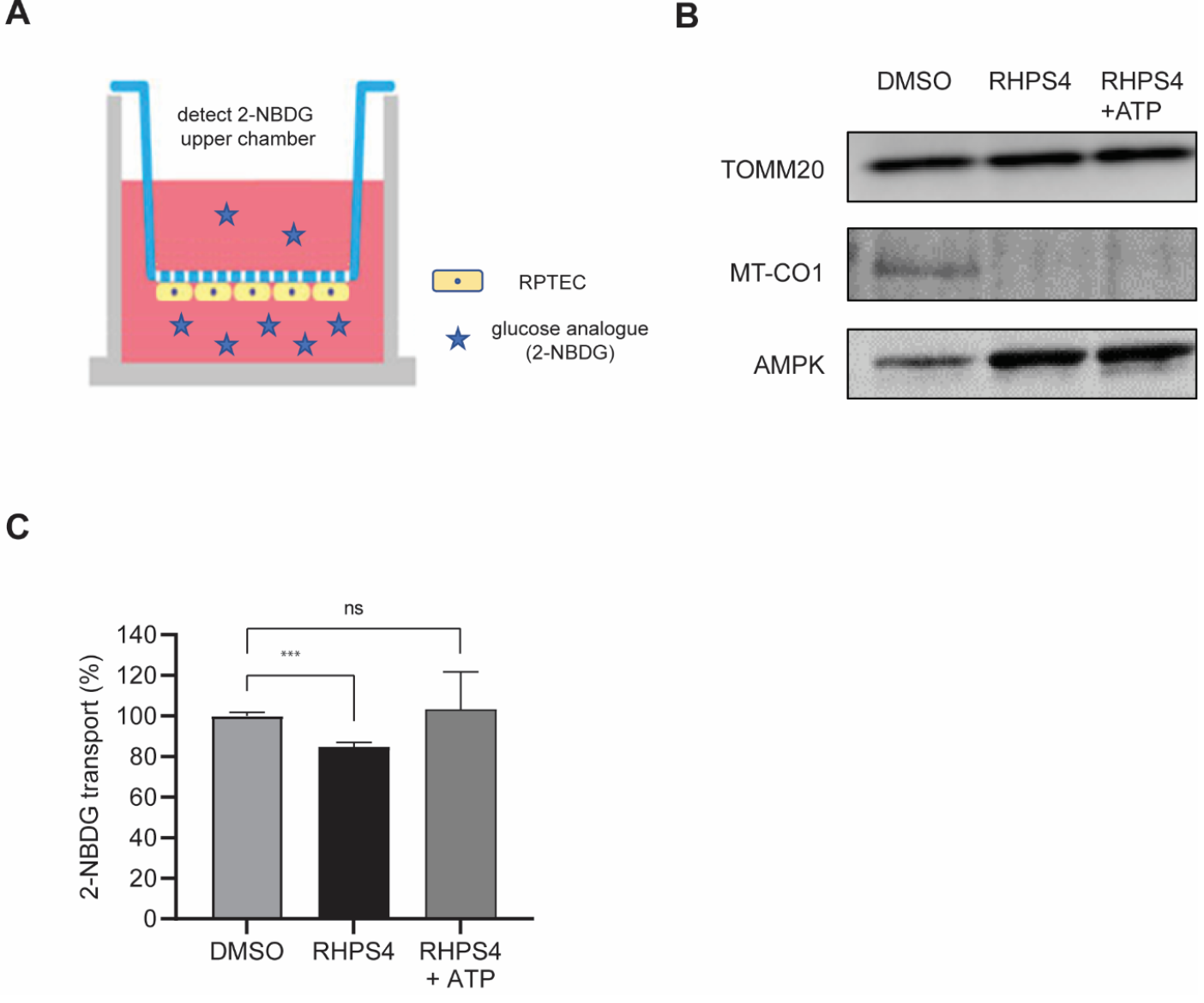
mtRNAP pausing impairs renal proximal tubule function. (A) Schematic of renal proximal tubule culture system, where cells are maintained on the bottom of the permeable membrane. The lower chamber is apical to the cells and the upper chamber is basal. Fluorescent substrates are placed in the lower chamber and active transport measured by detection of fluorescence in the upper chamber. (B) Protein expression nuclear encoded TOMM20, mitochondria encoded CO-1 and AMPK before and after 72 hrs of RHPS4 treatment (N=1). (C) Active transport of glucose analogue 2-NBDG, expressed as a percentage of transport performed by DMSO-treated cells. (N=5; ***P<0.001, t-test, error bars=S.E.M).

Next, we carried out a transporter assay to assess how the G4 mediated mtRNAP pausing affects the function on the renal epithelial cells. We measured impedance by transepithelial electrical resistance and observed a >5-fold (N=3, P<0.0001; one-way ANOVA) increase in the integrity of the epithelial barrier over 4 weeks in culture (Supplemental Figure 4A). To ask if the epithelial cells are a barrier to passive diffusion, we added fluorescently labelled dextran to the bottom chamber of the transwell and measured fluorescence in the upper chamber. We found 50-fold lower (N=3, P<0.0001; t-test) fluorescence in the upper chamber indicating the proximal tubule cells form an intact epithelial barrier (Supplemental Figure 4B). We then treated cells with the glucose analogue 2-NBDG in the presence of RHPS4 or vehicle. The results showed that with a decrease in ATP generation there was a significant (P<0.001) decrease in 2-NBDG transport (Figure 4D). The impairment in transport activity was fully rescued with the addition of ATP following RHPS4 treatment, showing that glucose transporters are not inhibited by RHPS4, but lack the energy for active transport. Together, this indicates that altered G4-mediated pausing of mtRNAP results in decreased ATP generation and is sufficient to impair the function of renal proximal tubule cells.

## Discussion

Understanding the effect of cis-elements on RNA polymerase pausing is essential to understanding transcriptional regulation. Here, by taking advantage of the unique architecture of the mitochondrial genome we could assess the relationship between guanine quadruplexes and mtRNAP pausing. Mitochondrial genes are encoded as polycistronic transcripts with the 37 genes synthesized from only two major promoters(1). The genome does not associate with histones, and the pausing complexes NELF and DSIF are not known to localize to mitochondria. We found mtRNAP paused at hundreds of locations where the polymerase has transcribed through guanine-rich regions where G4s form. We confirmed the formation of G4 both through computational and experimental approaches.

In the gene body of the mitochondrial polycistron we find that mtRNAP pauses after transcribing through G-rich regions where G4 form. Because the two mitochondrial polycistrons cover nearly the entirety of the mitochondrial genome we could correlate the location of mtRNAP pausing and the underlying sequence elements on both strands. We observed the mtRNAP paused after synthesis of a guanine-rich RNA. This implies G4s in the gene body do not act by forming a physical barrier in front of the polymerase, but instead may induce conformational changes in the elongation complex. Cramer and colleagues showed G4 formation in nascent RNA can destabilize the mtRNAP elongation complex (28) and this may be sufficient to cause the observed transcriptional pauses during transcription elongation.

Our results link the stabilization of G4s in the presence of RHPS4 to decreased mtRNAP transcription, lower mtRNAP gene expression, and impaired mitochondrial function. Decreased mitochondrial gene expression due to RHPS4 has been observed previously and was proposed to be the result of a defect in mtRNAP transcription(26) though the mechanism was not clear. Our findings here provide a mechanism for that observation: stable G4s result in more mtRNAP pausing and consequently decreased transcription. This has implications for cell types with high metabolic demands, such as renal epithelia, that depend on mitochondrial energy production and suggests that dynamic tuning of nucleic acid secondary structure can regulate mitochondrial transcription.

## Data availability

The deep sequencing data reported in this paper have been deposited in the NCBI sequence read archive PRJNA1028619

## Acknowledgments

We thank Drs. Robert Petrovich and Randy Bledsoe of the Structural Biology Core, Dr. Geoffrey Mueller of the NIEHS NMR Research Core Facility, Rickie Fannin of the NIEHS Microarray Core Facility, and Dr. Jeff Tucker and Erica Scappini of the NIEHS Fluorescence Microscopy and Imaging Core.

