## Supplemental Data for "Guanine quadruplexes mediate mitochondrial RNA polymerase pausing"

Supplemental Figures 1-4
Supplemental Table 1

Supplemental Figure 1

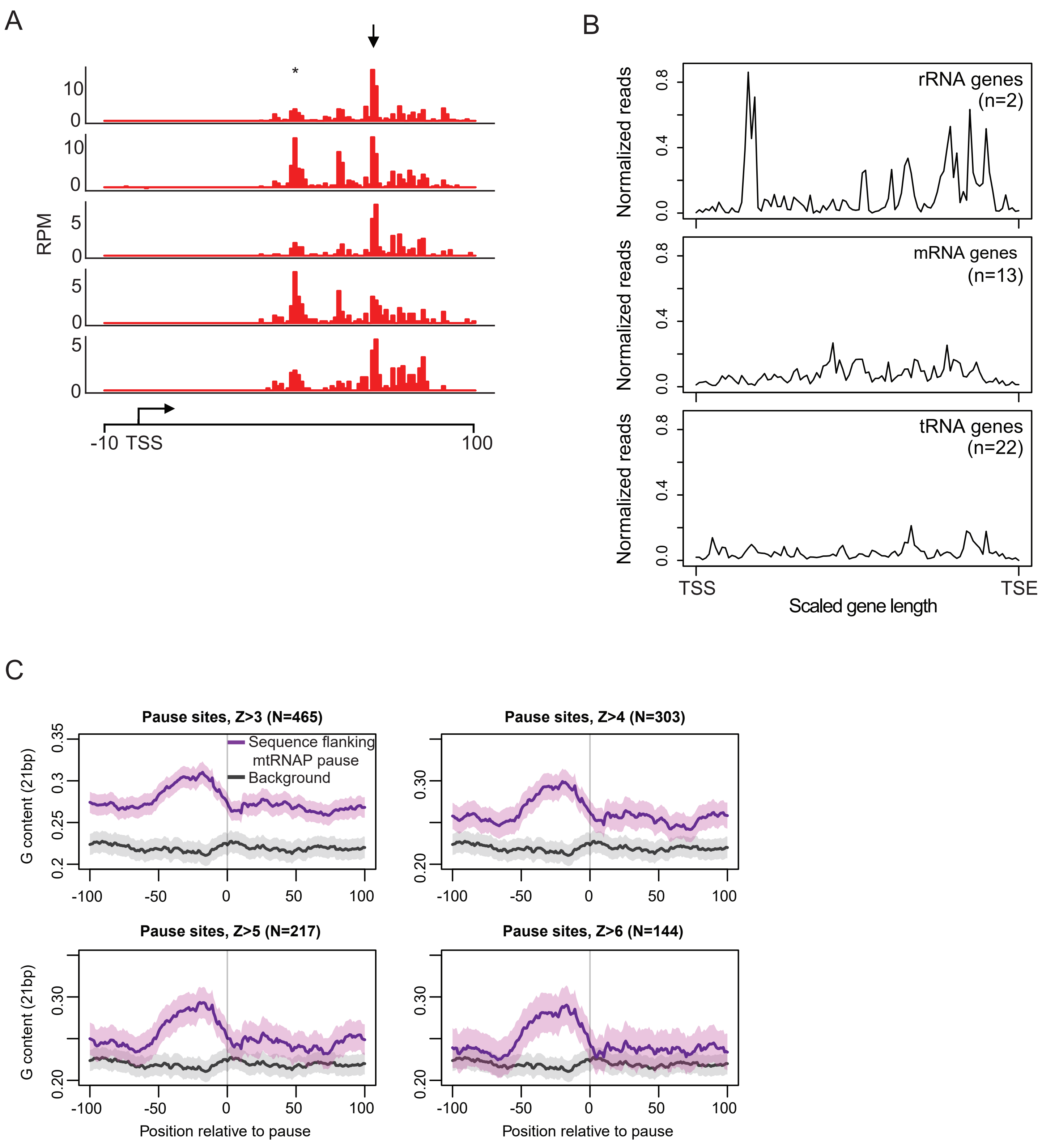


**Supplemental Figure 1. mtRNAP pauses proximal to the light strand promoter** (A) PRO-seq coverage on the light strand in primary fibroblasts (N=5). The TSS (arrow) and known site of mtRNAP pausing (*) are indicated. (B) Metagene plots of the mtRNAP distribution at rRNA, mRNA, and tRNA genes indicate mtRNAP pausing is not enrichment near the beginning or end of transcripts. (C) Guanine content near pause sites at increasingly stringent Z-score threshold to call pause sites, as compared to background sequence where mtRNAP does not pause (black).

Supplemental Figure 2

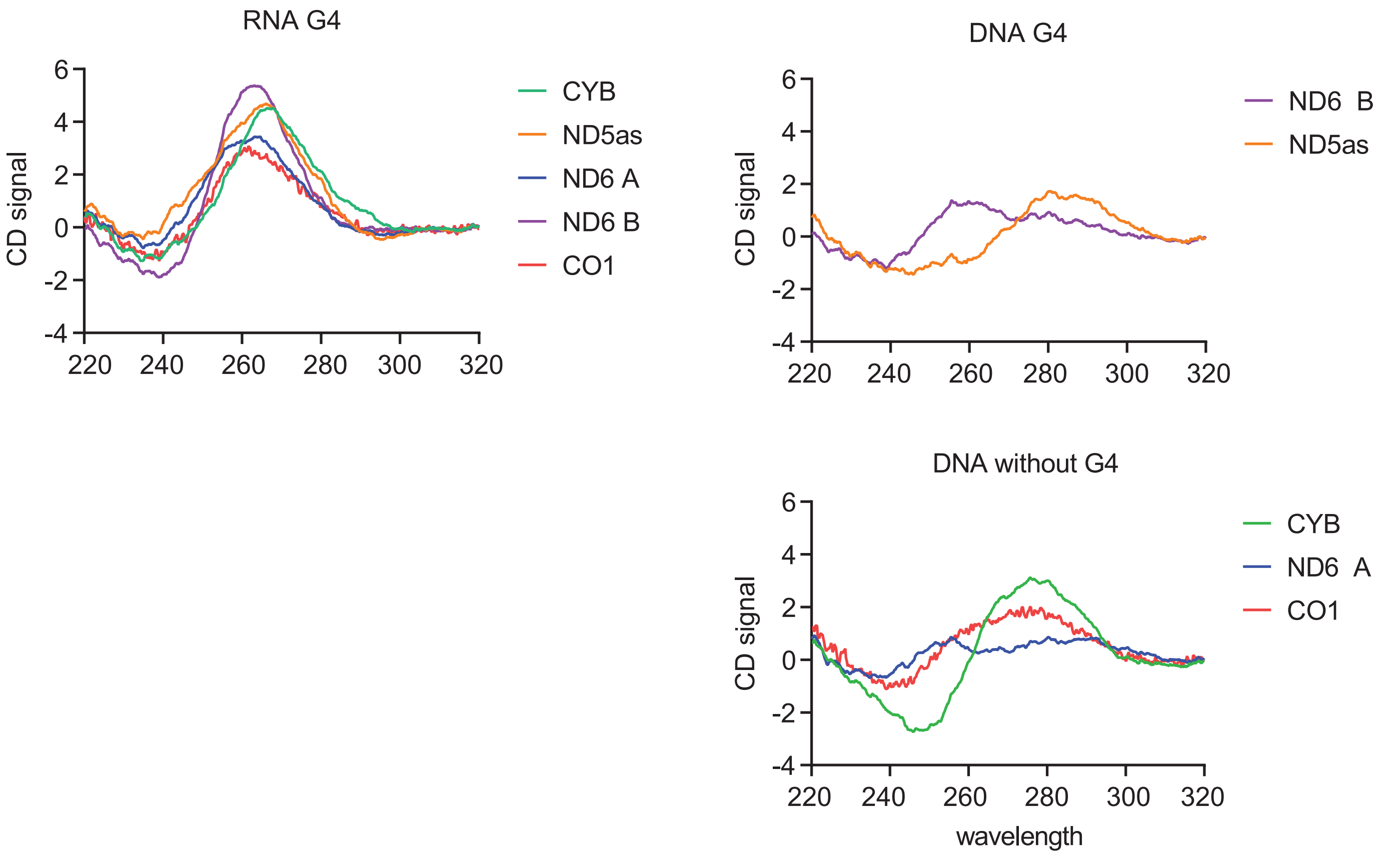


**Supplemental Figure 2. G4 formation is more stable in RNA.** Circular dichroism spectra of RNA or DNA oligos. Spectra from RNA oligos are consistent with parallel quadruplexes. DNA oligos are heterogenous with parallel or antiparallel quadruplexes (upper plot) or unfolded molecules (lower plot).

Supplemental Figure 3

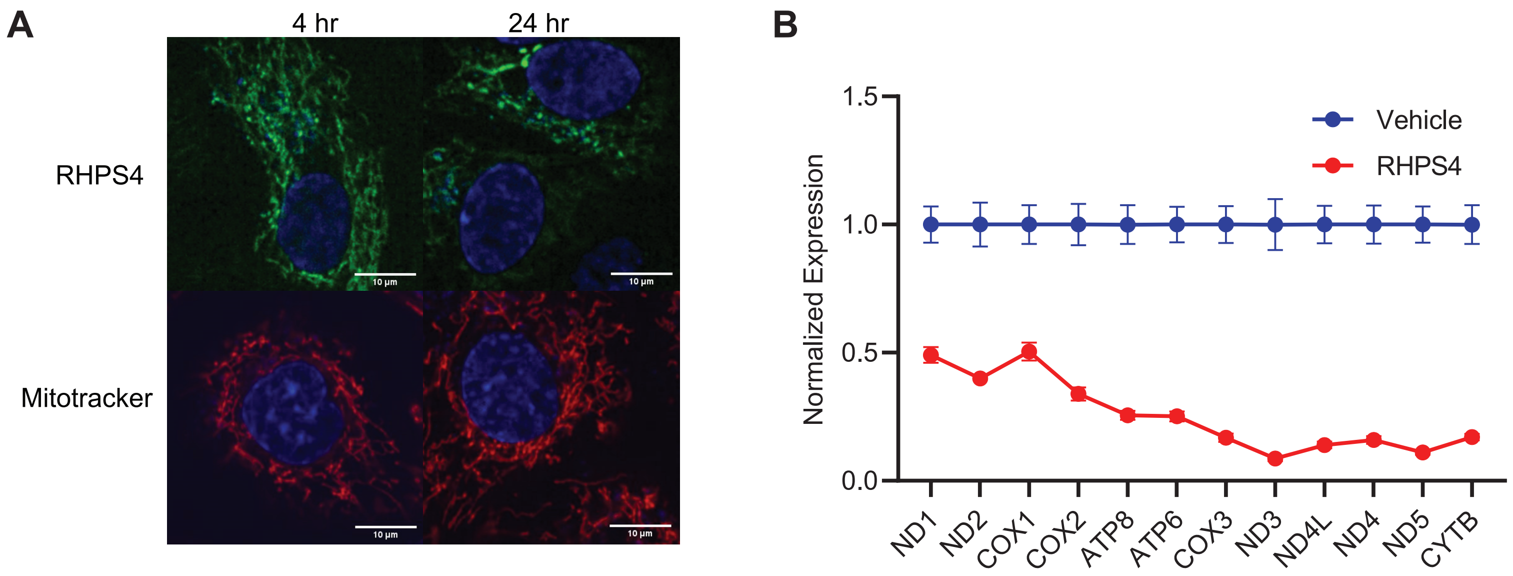


**Supplemental Figure 3. mtRNAP transcription is decreased when G4 are stabilized.** (A) Confocal image showing RHPS4 (green) uptake into mitochondria following 24 hours of drug treatment. (B) Mitochondrial gene expression in RPTEC before and 24 hours after RHPS4 treatment normalized to vehicle treated cells. Gene expression is in arbitrary units (N=3; $P<0.0001$, one-sided ANOVA, error bars=S.E.M.).

Supplemental Figure 4

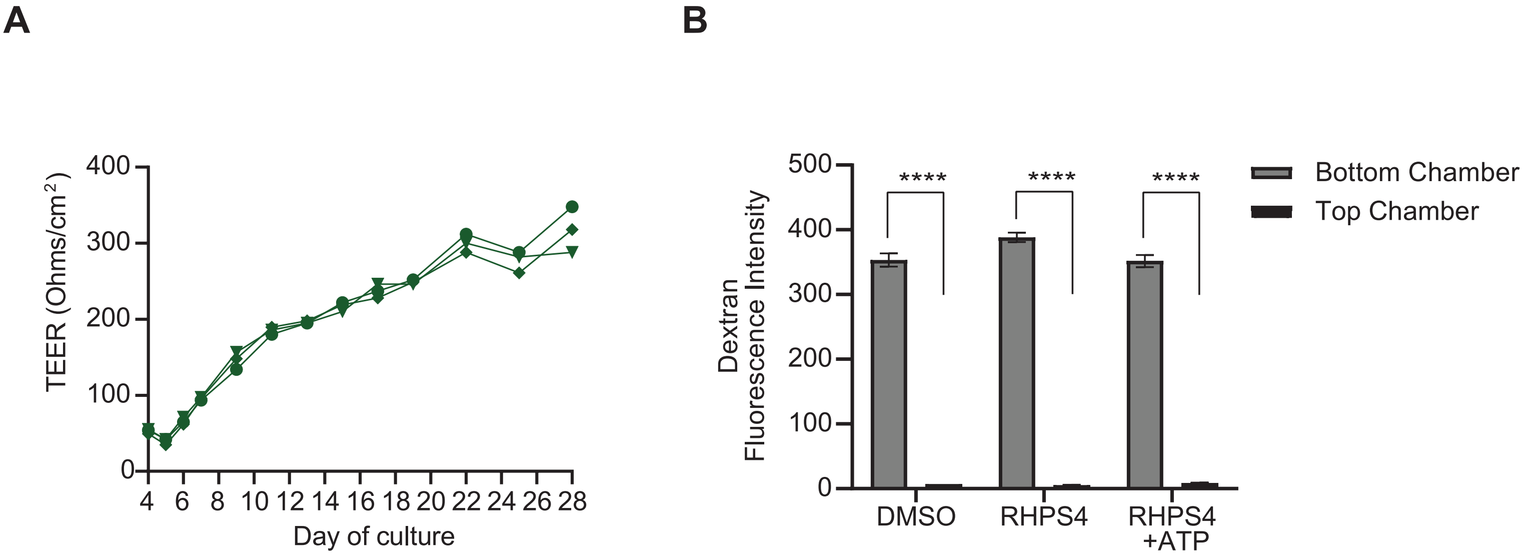


**Supplemental Figure 4. RPTEC monolayers form a barrier to diffusion.** (A) 5-fold increase in transepithelial electrical resistance (TEER) measurements over 30 days of culture (N=3, $P<0.0001$; one-way ANOVA). (B) RPTEC monolayers prevent diffusion of fluorescently labelled dextran from the bottom chamber to the top chamber in a transwell.

**Supplemental Table 1: Oligonucleotides**

| Oligo Name: | Sequence: |
|---|---|
| CO1_ChIP_F | TACCTCCCTCTCTCCTACTCCT |
| CO1_ChIP_R | GGCGTTTGGTATTGGGTTATGG |
| CO1_RNA | UGGAGGCCGGAGCAGGAACAGGU |
| CO1_RNA_mut | UUUAUUCCUUAGCAUUAACAUUU |
| CO1_DNA | TGGAGGCCGGAGCAGGAACAGGT |
| CYB_RNA | CGGGCGAGGCCUAUAUUACGGA |
| CYB_DNA | CGGGCGAGGCCTATATTACGGA |
| ND5AS_RNA | AGGCCUAGAUAGGGGAUUGUGCGGUGUGUGAUG |
| ND5AS_DNA | AGGCCTAGATAGGGGATTGTGCGGTGTGTGATG |
| ND6A_RNA | UGGAGGUAGGAUUGGUGCUGUGGGU |
| ND6A_DNA | TGGAGGTAGGATTGGTGCTGTGGGT |
| ND6B_RNA | UGAUGGGGUGGUGGUUGUGGU |
| ND6B_DNA | TGATGGGGTGGTGGTTGTGGT |
